# Src acts with WNT/FGFRL signaling to pattern the planarian anteroposterior axis

**DOI:** 10.1101/2021.08.18.456902

**Authors:** Nicolle A. Bonar, David I. Gittin, Christian P. Petersen

## Abstract

Tissue identity determination is critical for regeneration, and the planarian anteroposterior (AP) axis uses positional control genes expressed from bodywall muscle to determine body regionalization. Canonical Wnt signaling establishes anterior versus posterior pole identities through *notum* and *wnt1* signaling, and two Wnt/FGFRL signaling pathways control head and trunk domains, but their downstream signaling mechanisms are not fully understood. Here we identify a planarian Src homolog that restricts head and trunk identities to anterior positions. *src-1(RNAi)* animals formed enlarged brains and ectopic eyes and also duplicated trunk tissue, similar to a combination of Wnt/FGFRL RNAi phenotypes. *src-1* was required for establishing territories of positional control gene expression, indicating it acts at an upstream step in patterning the AP axis. Double RNAi experiments and eye regeneration assays suggest *src-1* can act in parallel to at least some Wnt and FGFRL factors. Co-inhibition of *src-1* with other posterior-promoting factors led to dramatic patterning changes and a reprogramming of Wnt/FGFRLs into controlling new positional outputs. These results identify *src-1* as a factor that promotes robustness of the AP positional system that instructs appropriate regeneration.

**Highlights:**

1. *Src-1* suppresses head and trunk identity
2. *Src-1* can regulate positional control gene domains
3. *Src-1* likely acts independently of *notum*/Wnt and FGFRL signals
4. *Src-1* inhibition broadly sensitizes animals to AP pattern disruption

## Introduction

Robust pattern control is fundamental to the process of regeneration (Wolpert, 1969). Animals must be able to re-establish tissue identity and proper polarity after injury for regeneration to proceed normally. Furthermore, regardless of regeneration abilities, many animals must also maintain regional identity throughout adult life as they replace and specify new cells to replenish old tissue. Planarians present a powerful system for studying these patterning control mechanisms, as they possess a remarkable ability to regenerate any missing body part and are in a state of constant cellular turnover to replace aged tissues (Elliott and Sanchez Alvarado, 2013; Reddien, 2018; Rink, 2018). Planarian regeneration abilities extend from a population of pluripotent stem cells, termed neoblasts, which continuously produce all adult cell types (Wagner et al., 2011; Zeng et al., 2018). Planarian muscle cells harbor positional information used in controlling neoblast differentiation and targeting through expression of regionalization determinants termed positional control genes (PCGs) (Witchley et al., 2013; Scimone et al., 2017). PCGs include signaling molecules in the Wnt, FGF, and BMP pathways that control tissue identity along the anteroposterior (AP) from head to tail, the dorsoventral (DV) axis from back to belly and the mediolateral axis. These factors are expressed in regional territories in uninjured animals that are reset during the regeneration process, and their inhibition leads to mispatterning phenotypes. However, the signaling mechanisms controlling positional information domains in muscle are not yet fully understood.

Significant progress has been made in understanding the regeneration of the planarian AP axis, which is driven by variants of Wnt signaling. Several of the nine planarian Wnt genes are expressed in overlapping domains from the posterior, while several Wnt inhibitors demarcate nested anterior domains. In recent years, the functions of many of these factors have been elucidated. A canonical β-catenin-dependent Wnt signaling pathway controls the head-versus-tail identity of blastemas after transverse amputation. Downregulation of Wnt pathway components β*-catenin-1*, *wnt1, Evi/wntless, Dvl-1/2* or *teashirt* causes regeneration of ectopic heads (Petersen and Reddien, 2008; Petersen and Reddien, 2009; Gurley et al., 2010; Iglesias et al., 2011; Owen et al., 2015; Reuter et al., 2015) whereas up-regulation of canonical Wnt signaling via RNAi inhibition of the Wnt negative regulators *notum* and *APC* causes the regeneration of ectopic tails (Gurley et al., 2008; Petersen and Reddien, 2011). *wnt1* and *notum* are both transcriptionally induced by injury where they likely participate in the control of polarization or orientation of the outgrowing blastemal tissue. An activin-dependent process restricts the initial 6-18 hours of *notum* expression to anterior-facing wounds, resulting in a low Wnt environment that leads to head regeneration (Cloutier et al., 2021). At later times in regeneration (by 24-72hours) and throughout homeostasis, stem cell-dependent processes (Hayashi et al., 2011; Currie and Pearson, 2013; Marz et al., 2013; Scimone et al., 2014; Vasquez-Doorman and Petersen, 2014; Vogg et al., 2014; Tejada-Romero et al., 2015; Schad and Petersen, 2020) generate cells expressing *wnt1* and *notum* in muscle cells at the posterior and anterior midline termini respectively (termed poles) where they may function to control region-specific patterning or act at the tip of a hierarchy of AP regulatory factors (Adell et al., 2009; Petersen and Reddien, 2009; Gurley et al., 2010; Stuckemann et al., 2017; Schad and Petersen, 2020).

Other Wnt-dependent pathways may function downstream or in parallel to pole identity and tissue polarization. *wnt11-6* (also known as *wntA*) and associated factors limit the regionalization of head tissue. Inhibition of *wnt11-6* or the *fzd5/8-4* Wnt receptor causes posterior expansion of the brain and the formation of ectopic posterior eyes (Kobayashi et al., 2007; Adell et al., 2009; Hill and Petersen, 2015; Scimone et al., 2016). Similarly, RNAi of *nou-darake (ndk),* a member of the FGFR-like (FGFRL) family of putative FGF decoy receptors, also results in a brain expansion phenotype along with ectopic posterior eyes (Cebria et al., 2002). The Wnt inhibitor *notum* also acts oppositely in the head regionalization pathway and independent of its roles in *wnt1/*polarity signaling. *notum(RNAi)* decapitated animals that succeed in regenerating a head form a miniaturized brain with elongated eyes, and also *notum(RNAi)* regenerating head fragments attain a reduced sized brain and form an ectopic set of anterior photoreceptors (Hill and Petersen, 2015). *notum* likely acts mainly through *wnt11-6* for anterior patterning because co-inhibition of *wnt11-6* suppresses the *notum(RNAi)* phenotypes of small brain and ectopic anterior photoreceptors, whereas co-inhibition of *wnt1* does not modify these phenotypes (Hill and Petersen, 2015; Atabay et al., 2018; Hill and Petersen, 2018). The restricted anterior expression of *notum* also suggests that head patterning is accomplished in part by maintaining a low-Wnt environment in the far anterior. *fzd5/8-4* and *ndk* expression is also restricted to the anterior region, while *wnt11-6* expression is prominent in the posterior brain and is also present in bodywall muscle across much of the AP axis. These factors are expressed constitutively, and their inhibition in uninjured animals leads to mispatterning phenotypes similar to those in regenerating animals (Hill and Petersen, 2015). Therefore, planarians use ongoing Wnt/FGFRL positional information to maintain anterior regionalization.

A separate set of Wnt-related and FGFRL genes control trunk identity in planarians. Inhibition of *ndl-3* (a FGFRL protein), *ptk7* (a kinase-dead Wnt co-receptor), *wntP-2* (Wnt ligand also called *wnt11-5*) (Gurley et al., 2010), or *fzd1/2/7* (Wnt receptor) causes posterior trunk duplication, with animals forming secondary mouths and ectopic pharynges (Lander and Petersen, 2016; Scimone et al., 2016). Similar to the anterior signals discussed above, these trunk patterning factors are required homeostatically and are expressed regionally. *wntP-2* is expressed in an animal-wide posterior-to-anterior gradient, *ptk7* is expressed in a trunk centered gradient, and *ndl-3* is expressed in a prepharyngeal territory. The head and trunk Wnt/FGFRL systems appear to act distinctly, because inhibition of one system does not influence the other’s phenotypic output. Together, these findings suggest that a body-wide system of Wnt-FGFRL signaling conveys positional information needed for regeneration and homeostatic tissue maintenance. Additional factors have been identified such as *nr4a* and *pbx* which regulate patterning at the termini (Blassberg et al., 2013; Chen et al., 2013; Li et al., 2019), *prep* transcription factor which regulates the anterior (Felix and Aboobaker, 2010), and *sp5* transcription factor which regulates territory within the tail (Tewari et al., 2019). In addition, Wnt and Activin signaling regulate fissioning behavior as well as the distribution of latent transverse regions prone to scission under pressure and which mark sites of future fission (Arnold et al., 2019). However, the downstream signaling factors important for body regionalization along the AP axis have not been fully resolved. In addition, it is not clear how signals from Wnt/FGFRL signaling along the AP axis relate to the canonical Wnt signaling used at the axis termini. Here we identify *src-1* as a global suppressor of anterior identities that can operate independently of pole formation. Our analysis indicates that *src-1* likely acts in parallel or downstream of pathways involving the Wnt and FGFRL factors to restrict anterior tissue identities in planarians.

## Results

### Planarian *src-1* Suppresses Head and Trunk Identity

To identify new regulators of regeneration patterning in planarians, we conducted an RNAi screen of 175 genes enriched for intracellular and receptor kinase activity (Table S1). Inhibition of 58 of these genes caused regeneration defects spanning from reduced blastema formation (21 genes), to aberrant photoreceptor formation (16 genes), to impaired movement behavior (5 genes). From this set, we identified a rare phenotype of ectopic eye formation in regeneration, due to inhibition of dd3147, a Src family homolog that we named *src-1* (Figure 1A, Supplementary Figure 1A). We isolated the clone and further analyzed this phenotype by staining to examine *src-1* requirements in patterning throughout the body. qPCR verified the effectiveness of the *src-1* dsRNA for *src-1* knockdown (Supplementary Figure 1B). Control animals regenerated two eyes as observed in live animals and measured by double fluorescence *in situ* hybridization (FISH) detecting both *opsin^+^* (eye photosensory neurons) and *tyrosinase* (eye pigment cup cells) (Figure 1A). By contrast, *src-1(RNAi)* animals formed ectopic posterior eyes in addition to their normal eyes (Figure 1A). Ectopic posterior photoreceptors formed in *src-1(RNAi)* regenerating fragments at a gradation of penetrance, highest in regenerating head fragments (89%, 180/203 animals), lower in regenerating trunk fragments (56%, 114/204 animals), and lowest in regenerating tail fragments (26%, 49/189). Therefore, *src-1* was most strongly required in anterior regions in order to prevent their posteriorization.

**Figure 1:**
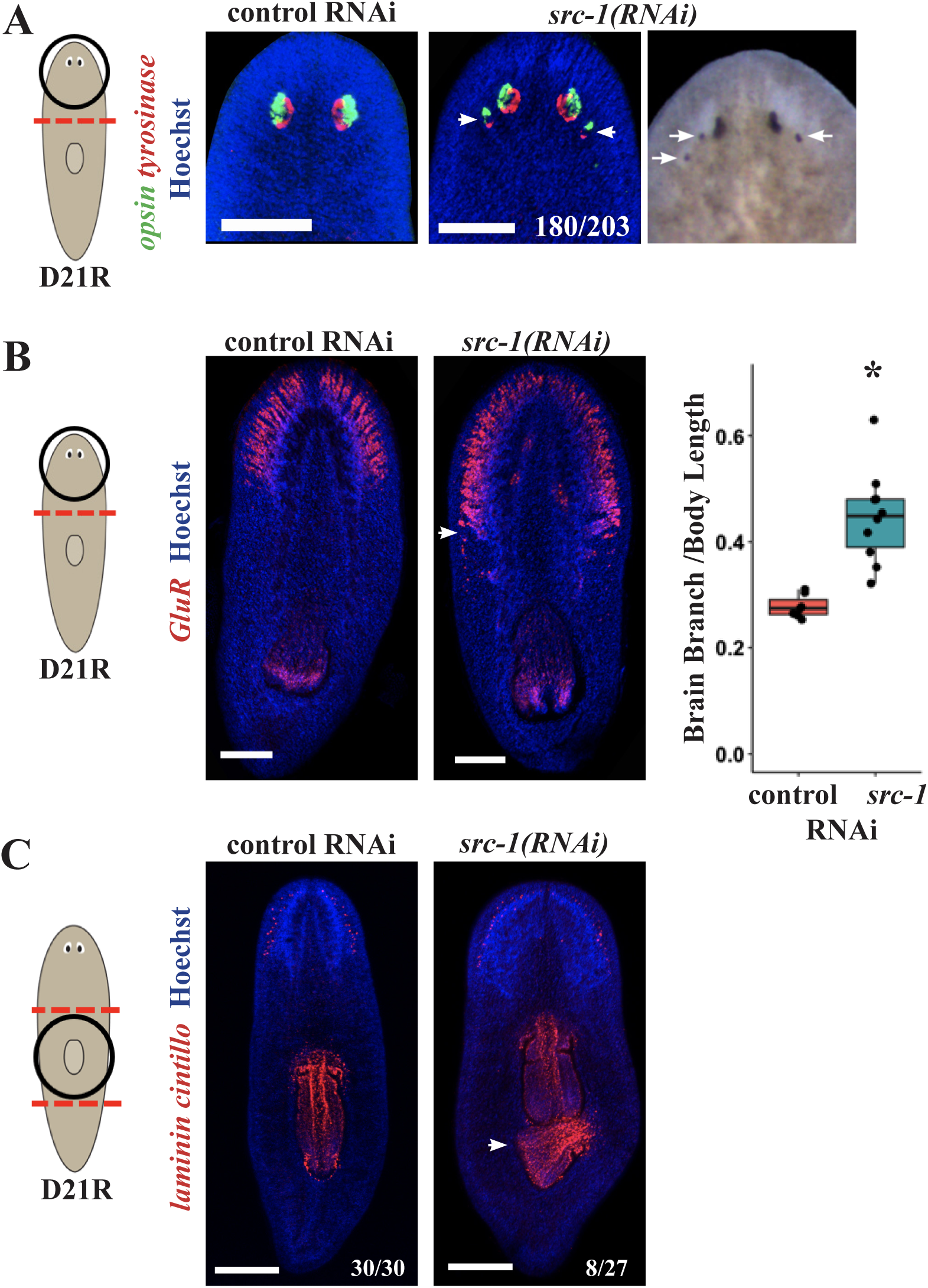
*src-1* restricts head and trunk identity to anterior positions. (A) *src-1(RNAi)* animals undergoing tail regeneration formed ectopic posterior eyes expressing *opsin* and *tyrosinase.* Scale bars, 150 µm (B) *src-1(RNAi)* animals undergoing tail regeneration formed a larger brain as evident by *GluR* expression, a marker of planarian brain branches. Scale bars, 300µm. Right, Quantification of brain branch length by *GluR* expression as proportional to body length, * p<0.05 by two-tailed t-test). (C) Regenerating *src-1(RNAi)* trunk fragments formed a posterior secondary pharynx (2 of 10 animals) as marked by *laminin* expression and a larger brain (10 of 10 animals) as marked by *cintillo* expression. Scale bars, 300µm.

The *src-1(RNAi)* eye phenotype was reminiscent of phenotypes observed for *ndk*, *wnt11-6*, and *fzd5/8-4* RNAi which also resulted in the formation of a larger brain (Cebria et al., 2002; Hill and Petersen, 2015; Scimone et al., 2016). Therefore, we sought to determine whether *src-1(RNAi)* animals similarly formed a larger brain. We investigated the size of the brain in *src-1(RNAi)* animals by examining the expression of *gluR,* a marker of the lateral brain branches, and found that *src-1(RNAi)* animals indeed formed a larger brain than controls that was posteriorly expanded (Figure 1B). Thus, we conclude that *src-1* acts to suppress head identity in general, similar to *wnt11-6* or *ndk* factors.

Given *src-1’s* requirement in regionalizing head identity, we tested whether it acted specifically in this process versus more generally in other AP patterning. Several factors have been implicated in restricting trunk identity to a more anterior position, and these do not appear to influence head patterning: *fzd-1/2/7, ndl3, ptk7,* and *wntP-2* (Sureda-Gomez et al., 2015; Lander and Petersen, 2016; Scimone et al., 2016). Similar to inhibition of these regulators, *src-1(RNAi)* regenerating trunk fragments formed a secondary posterior pharynx (marked by *laminin* expression), at ∼30% penetrance (Figure 1C). Similar to observations for inhibition of *ndl-3*, *ptk7*, and *wntP-2*, ectopic pharynx phenotypes in *src-1(RNAi)* animals were only observed in regenerating trunk fragments that contained a pre-existing pharynx. Together, we conclude that *src-1* anteriorly limits both trunk and head domains and can act similarly to both the anterior and posterior Wnt/FGFRL systems.

### *src-1* is Broadly Expressed in Both Muscle and Non-Muscle Cells

We next investigated whether *src-1* could control positional control gene (PCG) expression as part of its function to regulate anterior patterning. We found *src-1* itself to be broadly expressed throughout the animal and not in a gradient-like fashion, differing from known PCGs (Supplementary Figure 2A). *src-1* expression was detected in both muscle and non-muscle cells (Supplementary Figure 2B) as measured by FISH and co-expression with the muscle marker *collagen*. These observations are consistent with single-cell RNA sequencing experiments which found *src-1* to be widely expressed in a wide variety of cell types, including muscle (Wurtzel et al., 2015) (Supplementary Figure 2C). Therefore, it is possible that *src-1* could act in muscle cells to regulate anterior identity in planarians, or alternatively influence patterning in some other way.

Muscle cells themselves are required for positional information, because selective depletion of muscle subtypes causes mis-patterning phenotypes. For example, *myoD* RNAi prevents stem cells from differentiating into longitudinal muscle cells, and as a result, animals fail to express *follistatin* and *notum* after injury, and anterior outgrowth fails. In addition, *nxk1-1* RNAi prevents differentiation of circular muscle fibers, leading to midline bifurcation and the formation of split heads in the anterior (Scimone et al., 2017). Therefore, we sought to determine whether *src-1* RNAi phenotypes could be explained by the absence of muscle cell bodies or their fiber projections. Immunostainings showed that the *src-1* RNAi animals possessed muscle fibers stained with the 6G10 antibody (Supplementary Figure 2D). In addition, muscle cell bodies labeled by the presence of *collagen* mRNA were also present in apparently normal distributions in regenerating *src-1(RNAi)* animals (Supplementary Figure 2E). This suggests that *src-1* regulates anterior patterning not through affecting muscle formation but instead by either changing the signaling within muscle or in other cell types.

### *src-1* can pattern the AP axis independently from pole identity

We next tested whether expansion of the head region in *src-1(RNAi)* animals may be a result of changes in the *notum-*expressing anterior pole region. We found *notum* to be asymmetrically expressed in *src-1(RNAi)* animals at 18 hours after amputation at similar levels to controls (Figure S3A), consistent with the observation that under these conditions *src-1(RNAi)* animals did not have impaired axis polarization. We also observed *notum* to be expressed anteriorly but localized more broadly at 72-hours post-amputation in *src-1* RNAi animals, suggestive of a delay in pole formation (Figure S3B). By 14-days after amputations, all *src-1(RNAi)* animals had succeeded in regenerating a *notum+* anterior pole which was mildly expanded laterally. *notum* is also expressed in anterior midline neurons of the brain (Hill and Petersen, 2015; Scimone et al., 2020), and in *src-1(RNAi)* animals brain-associated *notum* expression expanded posteriorly in concert with the expanded brain (Figure S3C-D).

In contrast to *notum*, *wnt1* is expressed at both the anterior- and posterior-facing wound sites after amputation and is required for the formation of the posterior pole in regenerating animals. Regenerating *src-1(RNAi)* animals had normal wound-induced *wnt1* expression at 18-hours post-amputation and formed a posterior pole by 72-hours post amputation (Figure 3A-B). Furthermore, after 14 days of regeneration or homeostatic inhibition, *src-1(RNAi)* animals had a normal posterior pole as compared to controls (marked by *wnt-1* expression) (Figure S3C-D). Thus, *src-1* inhibition did not strongly affect the establishment or maintenance of the posterior pole under conditions that could nonetheless lead to brain expansion. Posterior and anterior pole formation depends strongly on *β-catenin-1* and *APC* suggesting that *src-1* can act independently of these factors.

### *src-1* regulates expression of body-wide AP patterning factors

Given the expansion of the anterior pole in *src-1(RNAi)* animals, we next sought to examine whether the domains of anterior position control genes (PCGs) were similarly expanded in *src-1(RNAi)* animals. We examined the anterior PCGs *ndk* and *ndl-5* which are expressed in both brain and muscle cells. The occurrence of the brain expansion and ectopic eye phenotypes in head and trunk fragments suggested that *src-1* RNAi could likely alter pattern homeostatically, similar to inhibition of other PCGs (Lander and Petersen, 2016; Scimone et al., 2016). Both regenerating and uninjured *src-1(RNAi)* animals had expanded *ndk* and *ndl-5* domains that extended more posteriorly towards the pharynx than control animals (Figure 2, Figure S4). We next investigated possible *src-1-*dependent regulation of trunk patterning factors *ndl3, ptk7,* and *wntP-2* (Lander and Petersen, 2016; Scimone et al., 2016). *src-1* inhibition resulted in the reduction of the anterior boundary of *ndl3* and *ptk7* within the pre-pharyngeal region but did not impact the posterior boundary of these mRNAs (Figure 2). These observations suggest *src-1* acts to restrict the anterior domain in planarians and allows for the possibility that *src-1* could be activating *ndl-3* and *ptk-7* expression in order to control trunk identity. We then examined the effect of *src-1* inhibition on the trunk PCGs, *wntP-2*, expressed in a posterior-to-anterior gradient. *wntP-2* expression was unchanged in *src-1(RNAi)* uninjured animals (Figure 2). *axinB* is a negative regulator of Wnt/β-catenin signaling in planarians whose inhibition results in two-tailed planarians, and it is expressed similarly to *wntP-2* in a posterior-to-anterior gradient (Iglesias et al., 2011). Axins are feedback inhibitors of beta-catenin signaling, thus the expression of Axin marks locations of canonicl Wnt pathway activity. Unlike *β-catenin-1* RNAi (Lander and Petersen, 2016), *src-1* inhibition did not eliminate *axinB* expression, but because of the low expression of the *axinB* transcript in these experiments, we could not unambiguously rule out the possibility that *src-1* inhibition mildly modifies *axinB* expression in some way. However, this analysis suggests that *src-1* inhibition likely does not eliminate β-catenin signaling along the body axis, similar to prior observations made after *wntP-2* and *ptk7* RNAi (Figure 2)(Lander and Petersen, 2016).

**Figure 2:**
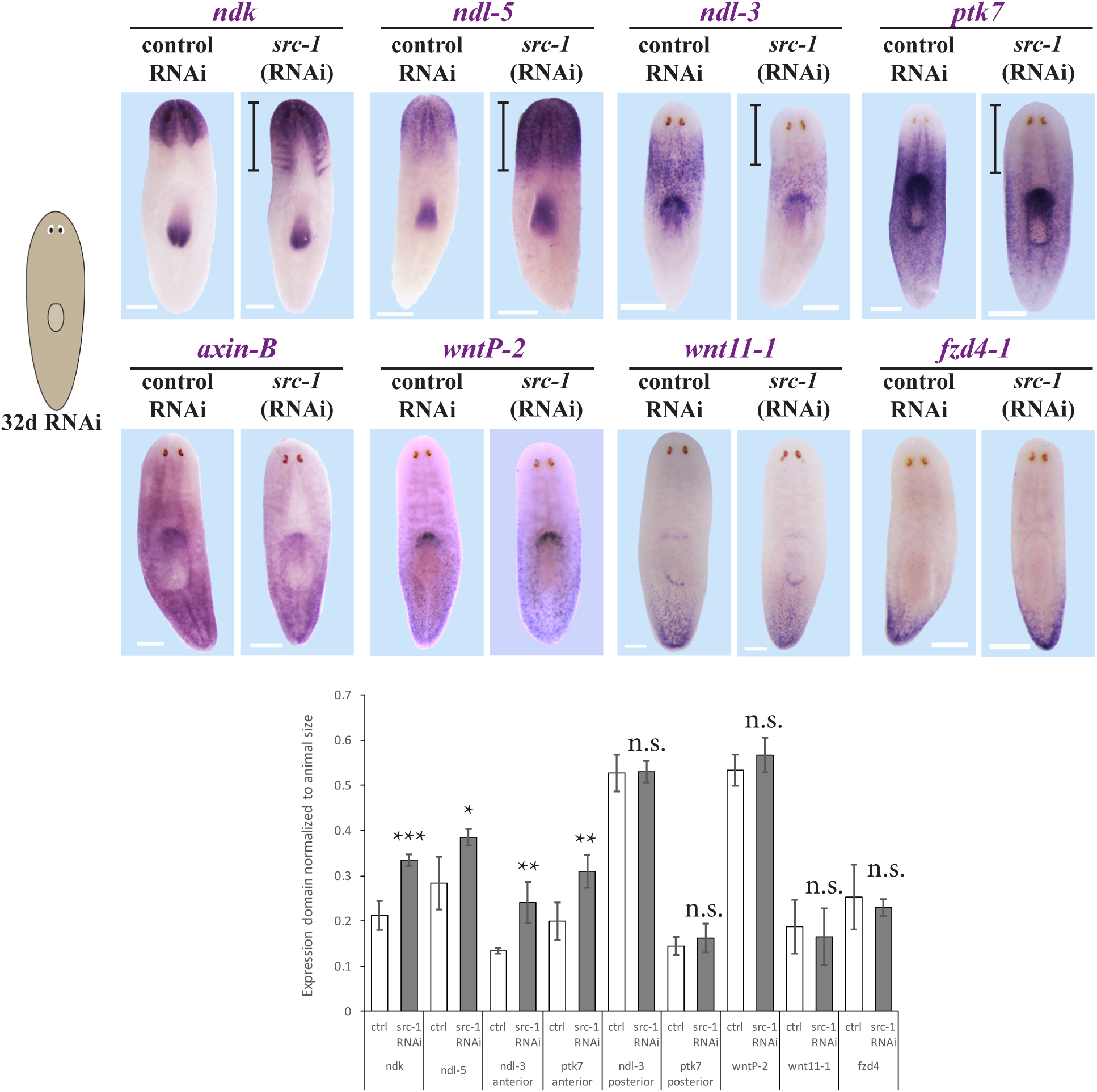
Anterior and central PCG domains are modified by *src-1* RNAi. Above, uninjured *src-1(RNAi)* or control RNAi animals stained for PCG domain expression by WISH as indicated after 32 days of gene inhibition. Bottom, quantifications of expression domains measured as a fraction of total animal length. Below, the PCG expression domain features were measured from the anterior animal tip (*ndk*, *ndl-5*, *ndl-3* anterior boundary, *ptk7* anterior boundary, *ndl-3* posterior boundary) or from the posterior tip (*ptk7* posterior boundary, *wntP-2*, *wnt11-1*, *fzd4*). At least 4 animals were used in each measurement. Asterisks mark significance from 2- tailed t-test, *p<0.05, ** p<0.01, *** p<0.001, n.s. p>0.05. *axinB* expression was continual across the axis and so could not be confidently scored in this way, and 4/4 animals appeared as shown. *src-1* inhibition caused a posterior shift to the anterior and central PCG domains (*ndk*, *ndl-5*, *ndl 3* anterior boundary, *ptk7* anterior boundary) and no significant change to posterior PCG domains.

Because *src-1(RNAi)* animals could regenerate an apparently normal posterior pole, we were interested to determine whether expression of other posterior PCGs would be affected after *src-1* RNAi. The expression of the posterior PCGs, *fzd-4,* and *wnt11-1* was unchanged in these *src-1(RNAi)* animals (Figure 2). Together, these observations point to a role for *src-1* in controlling anterior and central PCG expression domains.

### *src-1* likely acts independently of *notum/wnt11-6* in head patterning

Srcs are intracellular tyrosine kinase that can act as a signaling hub of multiple pathways and influence many cellular processes (Parsons and Parsons, 2004). Given that *src-1* inhibition shifts anterior PCG domains and results in brain expansion and posterior ectopic eye phenotypes reminiscent of *ndk* and *wnt11-6* RNAi (Cebria et al., 2002; Hill and Petersen, 2015), we sought to determine whether *src-1* might signal downstream of either factor. To begin to address this question, we designed epistasis experiments using double RNAi. *notum(RNAi)* head fragments form an ectopic set of eyes within the head tip anterior to the pre-existing photoreceptors, whereas *wnt11-6(RNAi)* head fragments form an ectopic set of eyes posterior to the pre-existing eyes. Concurrent inhibition of *notum* and *wnt11-6* has been shown to suppresses the anterior ectopic photoreceptor, arguing that *wnt11-6* likely acts downstream and oppositely to *notum* in head patterning (Hill and Petersen, 2015). We reasoned that if *src-1* acted primarily downstream of *wnt11-6,* and therefore of *notum* in the anterior, then dual inhibition of *notum* and *src-1* should produce the posterior ectopic eye and enlarged brain phenotypes seen in single inhibition of *src-1* while suppressing the *notum(RNAi)* anterior eye phenotypes. Instead, simultaneous inhibition of *notum* and *src-1* in amputated head fragments produced several different phenotypes: 24 of 42 animals exhibited a synthetic phenotype with both posterior and anterior photoreceptors, 9 of 42 animals had a *notum(RNAi)* phenotype with only anterior photoreceptors, 6 of 42 animals exhibited a *src-1(RNAi)* phenotype of only posterior photoreceptors, and 3 of 42 animals appeared normal (Figure 3A). The observation of a synthetic phenotype after inhibition of both *src-1* and *notum* at this frequency (in ∼50% of animals) indicates these factors can exert distinct influences and strongly suggests that *src-1* can act independently of *notum,* and therefore likely of *wnt11-6,* for controlling anterior identity. In support of this model, simultaneous inhibition of *notum* and *src-1* in amputated head fragments led to a brain size (as measured by *cintillo+* cell number) that was neither small like *notum(RNAi)* nor large like *src-1(RNAi)* but instead a size in between the two RNAi phenotypes (Figure 3B-C).

**Figure 3:**
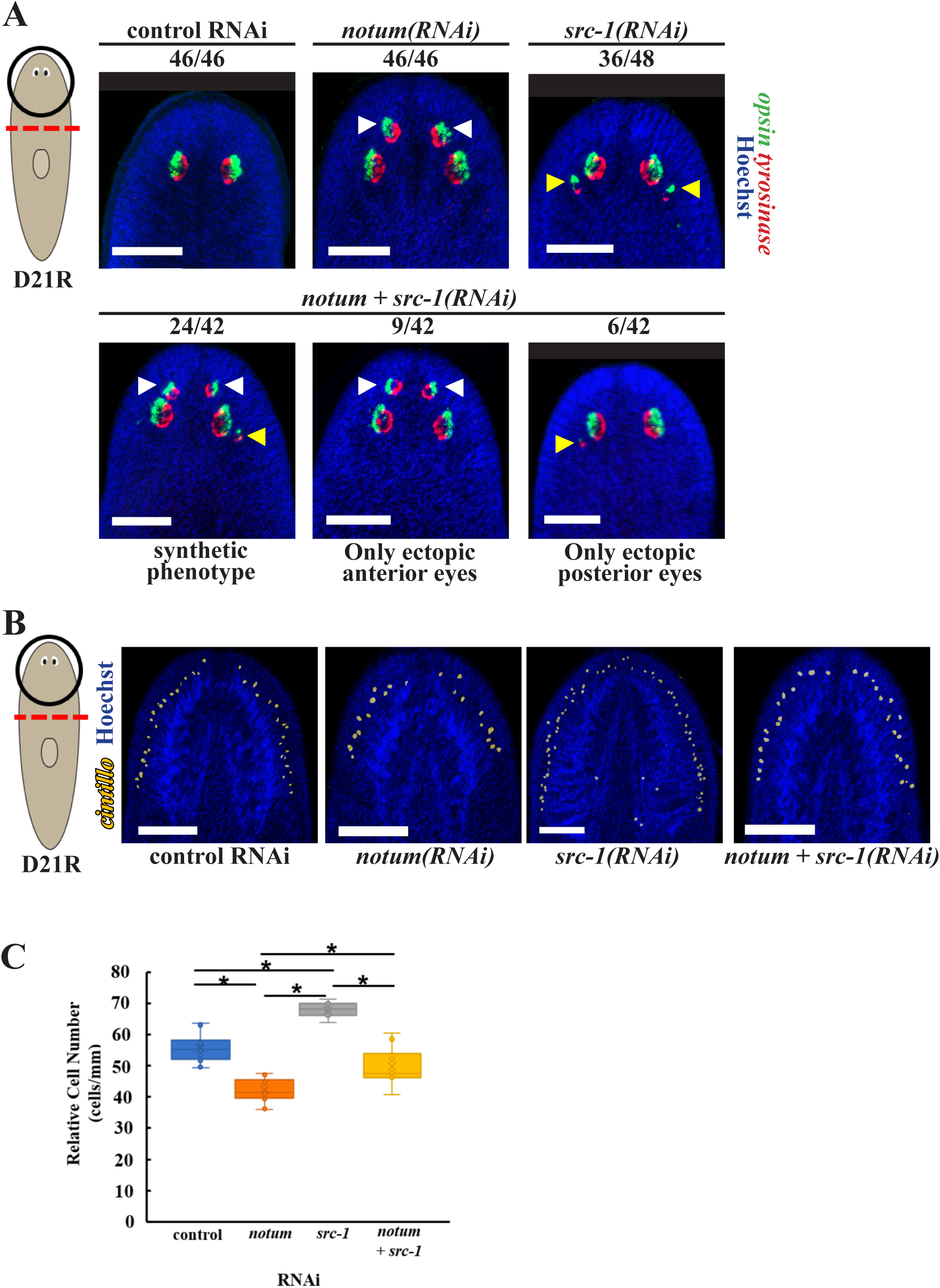
*notum* and *src-1* can act independently to determine eye placement. (A) FISH to detect expression of *opsin* (green), a marker of photoreceptor neurons, and *tyrosinase* (red), a marker of pigment cup cells, in control, *src-1, notum* and *src-1+notum(RNAi)* regenerating head fragments. Hoechst (blue) used as counterstain to detect nuclei. Ectopic eyes are marked by white arrows versus unmarked pre-existing eyes. *notum* RNAi caused formation of anterior ectopic eyes, and *src-1(RNAi)* caused the formation of posterior ectopic eyes, while simultaneous inhibition of *src-1 and notum(RNAi)* resulted in a synthetic phenotype in 24/42 animals with both anterior and posterior ectopic eyes. Scale bars, 150 microns. (B) FISH to detect expression of *cintillo* (red), a marker of chemosensory neurons, in control, *src-1, notum and src-1 + notum (RNAi)* regenerating head fragments. scale bars, 150 microns. (C) Quantification of *cintillo+* cell number normalized to animal size. *, p <0.05 from 2-tailed t-test. *notum* RNAi caused the regeneration with reduced numbers of *cintillo+* cells, and *src-1* RNAi caused formation of greater numbers of *cintillo+* cells, while simultaneous inhibition of *src-1* and *notum* resulted in an intermediate number of these cells.

### *src-1* and *wnt11-6* both act distinctly from *ndk* to define the location of eye regeneration

We next sought to test the possibility that *src-1* might transduce signaling through *ndk*. NDK/FGFRL receptors have ectodomains capable of binding FGFs but lack intracellular kinase domains to transduce signals, so have been proposed to act as FGF pathway decoy receptors. However, planarian FGF ligands have not been implicated in AP patterning, so it is unclear what other pathway components signal via planarian FGFRLs. The short intracellular domain of FGFRLs could be capable of recruiting other types of signaling models, so we considered the possibility that FGFRLs might signal through *src-1* by a close examination of the *ndk(RNAi)* versus *src-1(RNAi)* phenotypes. The *ndk* RNAi phenotype typically involves production of ectopic eyes at a more posterior location than *src-1* RNAi, giving some support for the distinct action of these factors.

We further probed the characteristics of the ectopic eyes in each RNAi condition, taking advantage of a newly identified distinction between *wnt11-6(RNAi)* and *ndk(RNAi)* conditions in controlling the location of eye regeneration after eye removal (Atabay et al., 2018; Hill and Petersen, 2018). Planarians under normal conditions can regenerate their eyes within ∼7 days after surgical removal. However, pattern disruption phenotypes resulting in ectopic eyes have distinct properties with respect to the location of regeneration after removal of ectopic versus pre-existing eyes. If the original, pre-existing photoreceptors in *wnt11-6*(RNAi) animals were surgically removed, they did not regenerate (11/11 animals). By contrast, when the ectopic photoreceptors of *wnt11-6(RNAi)* animals are surgically removed, new photoreceptors regenerated in that location a majority of the time (6/9 animals) (Figure 4). These results are consistent with prior studies (Atabay et al., 2018; Hill and Petersen, 2018) showing similar behavior for eyes in animals inhibited simultaneously for *wnt11-6* and *fzd5/8-4*. These results suggest that *wnt11-6* and *fzd5/8-4* signals control a target location for eye regeneration at a particular A/P position in the animal. By contrast, eye regeneration in *ndk(RNAi)* animals does not share this property, because in these animals removal of the pre-existing eyes still allowed for regeneration at that position most of the time (61%, 8/13 animals), while removal of ectopic photoreceptors did not lead to eye regeneration in a majority of animals (85%, 11/13 animals)(Figure 4)(Hill and Petersen, 2018). Therefore, *ndk* RNAi alters the overt pattern of animals without modifying the *wnt11-6-*dependent system directing the position of new eye regeneration.

**Figure 4:**
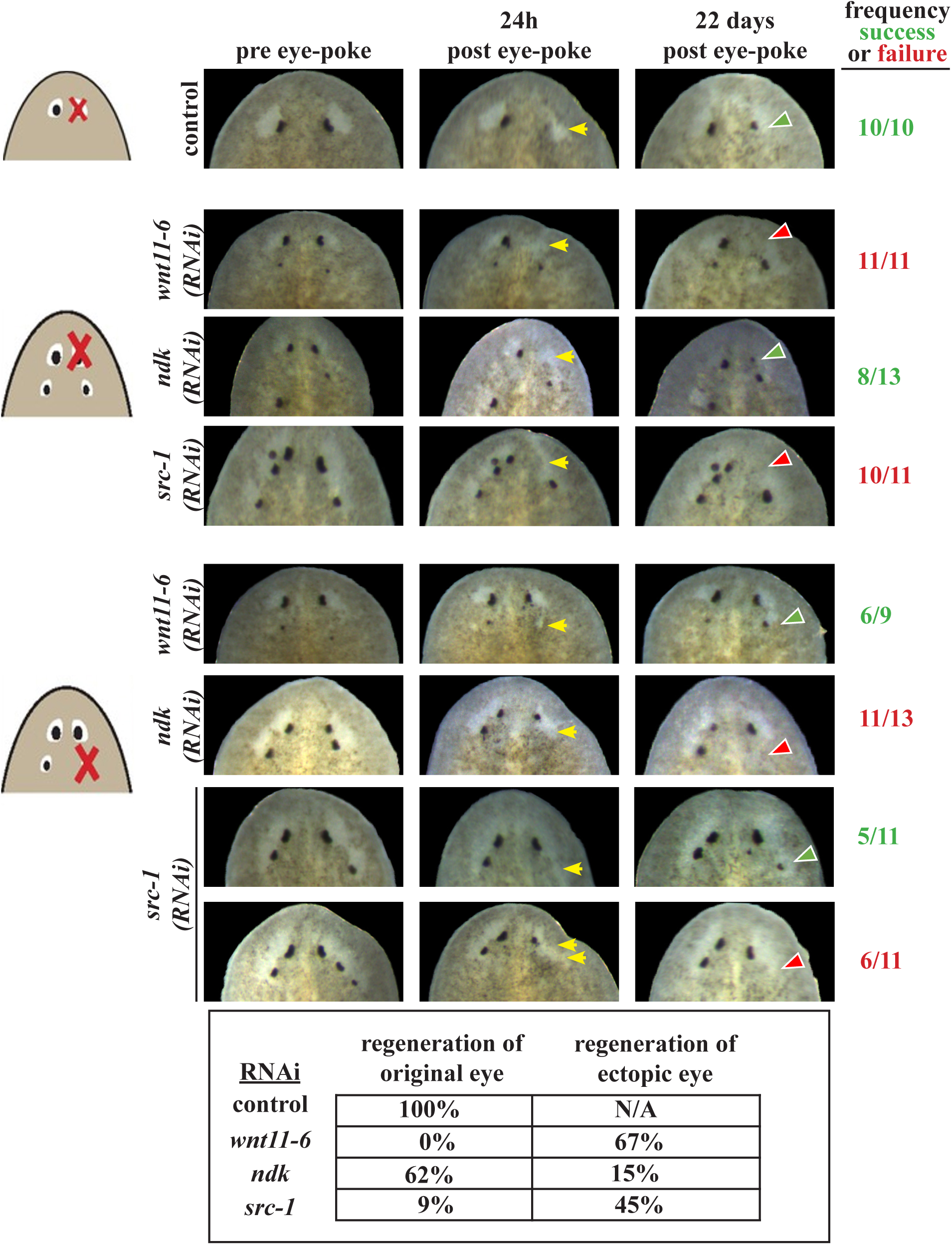
Inhibition of *src-1* or *wnt11-6* but not *ndk* alters the location of eye regeneration. Uninjured animals were fed the indicated dsRNA 12 times over 6 weeks and eye resection was then performed to remove either a pre-existing original eye or a supernumerary posterior eye in either *wnt11-6(RNAi)*, *ndk(RNAi)*, or *src-1(RNAi)* conditions. Animals were then imaged to verify eye removal (yellow arrows) and tracked individually as they attempted eye regeneration over the course of 22 days then scored for the successful (green arrows, green numbers scoring animals shown on right) or unsuccessful (red arrows, red numbers scoring animals shown on right) eye regeneration. Removal of the pre-existing eye resulted in successful eye regeneration in 8/13 *ndk(RNAi)* animals but in 0/11 *wnt11-6(RNAi)* animals and 1/11 *src-1(RNAi)* animals. By contrast, removal of the supernumerary eyes resulted in regeneration in 6/9 *wnt11-6(RNAi)* animals and 5/11 *src-1(RNAi)* animals but only 2/13 *ndk(RNAi)* animals. The bottom panel summarizes the frequency of regeneration from each condition and eye type. Therefore, either *wnt11-6* or *src-1* RNAi treatments shift the target location of eye regeneration to a more posterior position, while *ndk* RNAi did not as strongly cause this shift.

We reasoned that if *src-1* acted downstream of *ndk* to promote anterior identities, then as observed in *ndk(RNAi)*, the ectopic eyes in *src-1(RNAi)* animals would be incapable of regenerating but regeneration of pre-existing eyes would succeed. To test this model, we resected ectopic and pre-existing eyes from a cohort of homeostatic *src-1(RNAi)* animals then tracked each animal and eye profile individually over 22 days of recovery. Regeneration failed at the location of nearly all *src-1(RNAi)* transected pre-existing eyes (10 of 11 animals), and by contrast regeneration succeeded at the sites of transected ectopic eyes at a frequency (5 of 11 animals) close to that seen in *wnt11-6* RNAi (Figure 4). These results indicate that the site of eye regeneration is stably shifted after either *wnt11-6* or *src-1* RNAi but not after *ndk* RNAi. Therefore, *src-1* likely acts independently from *ndk.* Taken together, the double-RNAi and eye regeneration tests suggest that *wnt11-6, src-1,* and *ndk* likely control separate processes important in head patterning.

### *src-1* inhibition broadly sensitizes animals to AP pattern disruption

Given these findings of broadly parallel action, along with *src-1*’s role in maintaining anterior PCG domains, we reasoned that *src-1* inhibition might be capable of exerting a broader influence on positional signaling. To test this possibility, we carried out a series of double-RNAi experiments between *src-1* and other PCGs and examined effects on head and trunk patterning. We first simultaneously inhibited *src-1* with the head patterning factors *wnt11-6*, *ndk* and *fzd5/8-4*, each known to restrict eye cell number and brain size from more posterior regions, but which do not normally influence trunk identity in planarians (Scimone et al., 2016). Strikingly, when *src-1* was simultaneously inhibited with *wnt11-6*, *ndk*, or *fzd5/8-4*, animals had dramatically increased numbers of ectopic posterior eyes compared to any single gene RNAi conditions (Figure 5A). In addition, the double-RNAi animals had more severely expanded brains as measured by counting *cintillo+* chemosensory cells (Figure S5). Thus, inhibition of Src with anterior-specialized Wnt and FGFRL signals leads to enhanced anterior transformations.

**Figure 5:**
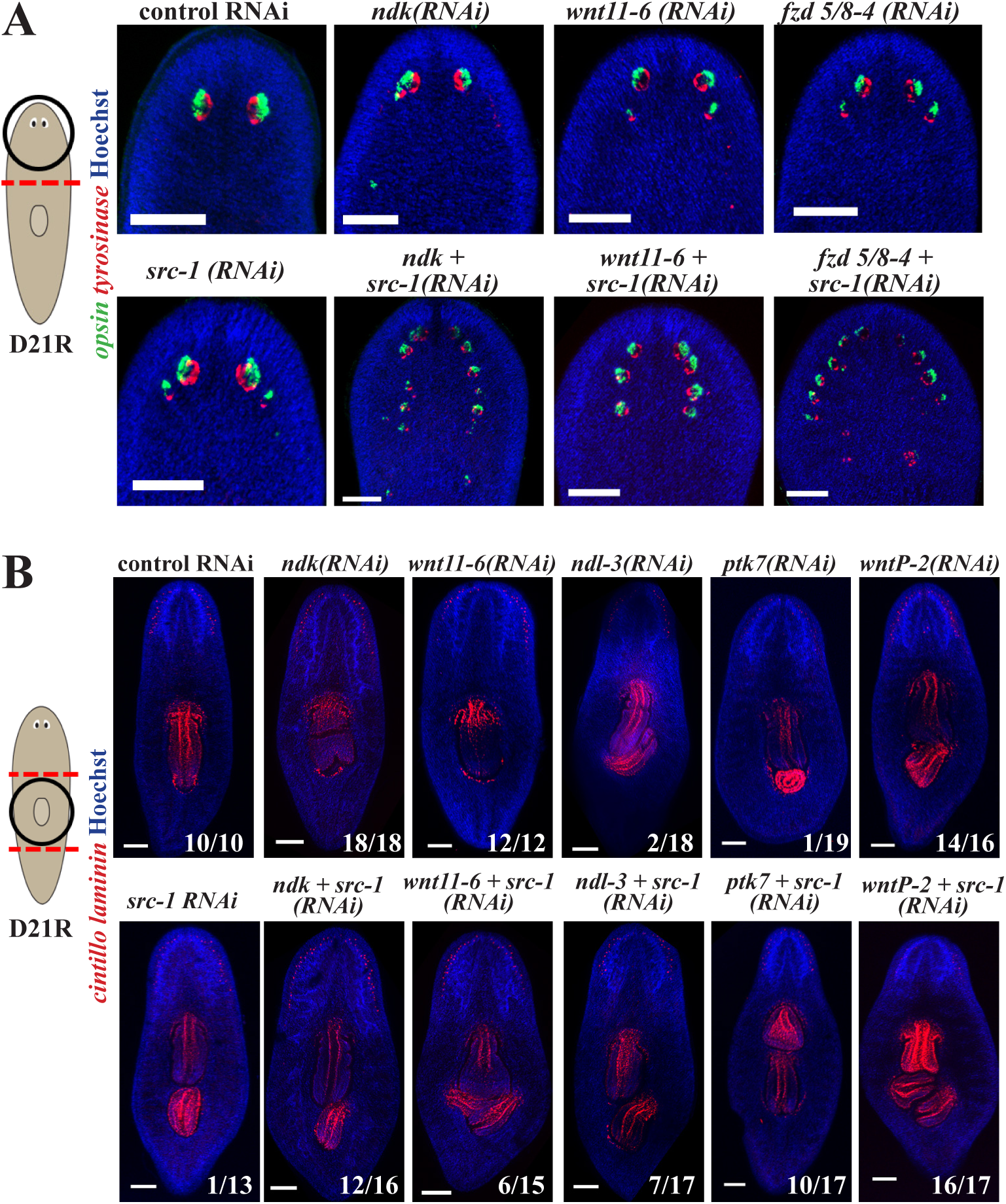
*src-1* inhibition sensitizes animals to AP pattern disruption and reprograms PCG activity. (A) FISH to detect expression of *opsin* (green), a marker of photoreceptor neurons, and *tyrosinase* (red), a marker of pigment cup cells, in head fragments at day 21 post amputation. Hoechst (blue) used as counterstain to detect nuclei. Simultaneous inhibition of *src-1* with *wnt11-6*, *ndk* or *fzd5/8-4* resulted in the formation of numerous ectopic eyes that extended posteriorly to a greater extent and number than in single-gene inhibitions. (B) Day-21 regenerating trunk fragments stained with *laminin* riboprobe to mark the pharynx (red, central), along with FISH of *cintillo* (red, anterior) marking chemosensory neurons. Simultaneous inhibition of *src-1* with *wnt11-6*, *ndk*, *ndl-3*, *ptk7*, or *wntP-2* resulted in the formation of ectopic posterior pharynges at a greater penetrance than each RNAi condition alone. Numbers indicate fraction of animals with either a single pharynx or ectopic pharynges as shown. Scale bars, 300um.

**Figure 6:**
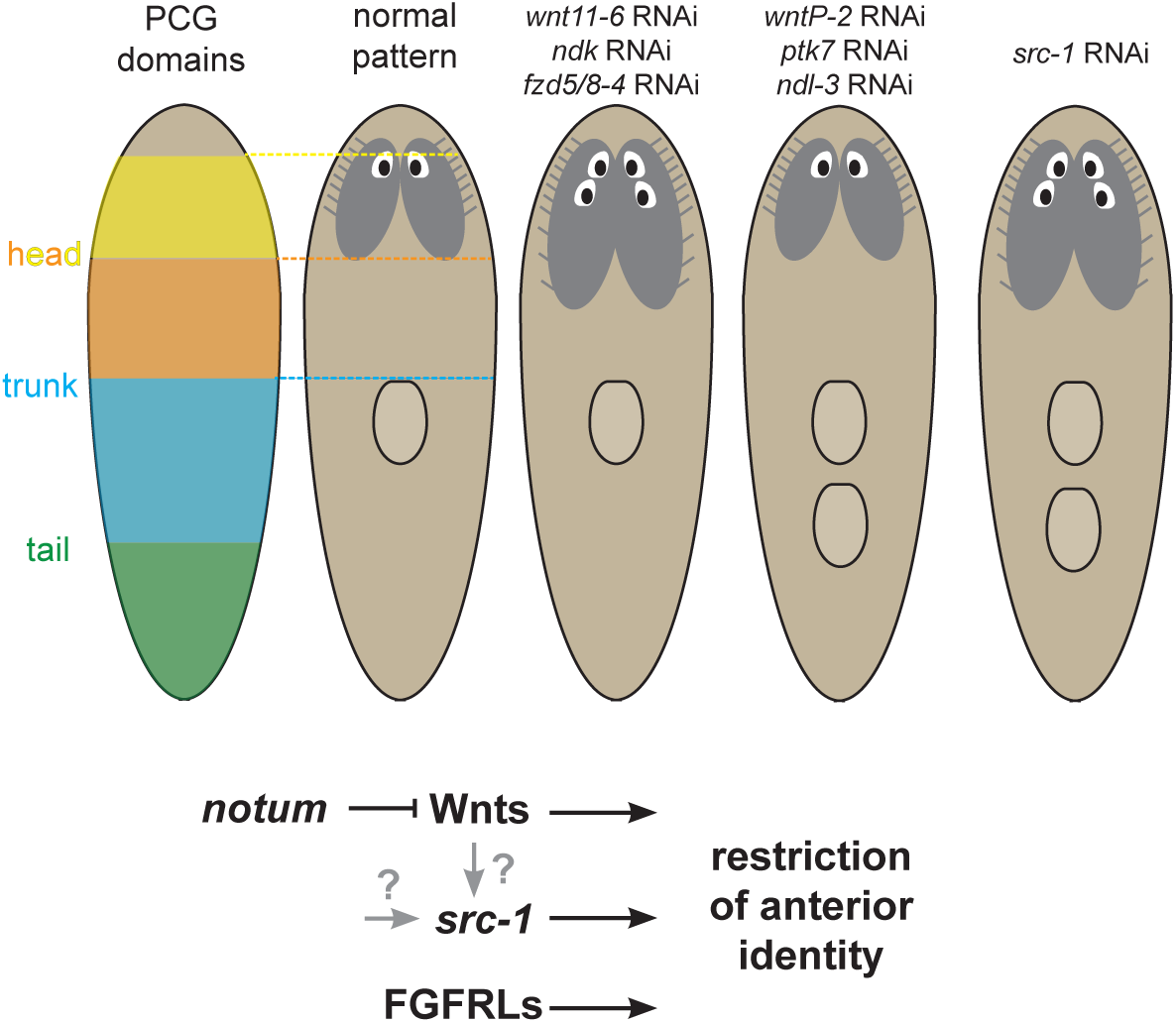
*src-1* acts with Wnt and FGFRLs to control AP axis identity. Model of *src-1* participating in A-P patterning along with Wnts and FGFRLs. Cartoons depict positional control gene domains from muscle that determine the normal animal pattern of eyes, brain and pharynx, as well as patterning phenotypes after inhibition of key factors. The synthetic phenotypes of *src-1* and *notum* RNAi suggest *src-1* can act independently of some Wnt signaling for control of AP identity. Likewise, the distinct effects of *src-1* RNAi and *ndk* RNAi on the location of eye regeneration after eye removal suggest that *ndk* likely does not act through *src-1*. However, it remains possible that *src-1* could act downstream of Wnts in a *notum*-independent process and/or downstream of other factors acting in parallel. *src-1* influences PCG domains and functions as a buffer to help define their territories and outputs, and thereby suppress anterior identity.

Next, we tested the effects of simultaneous *src-1* inhibition with the patterning factors known to restrict trunk but not head identity in planarians (Lander and Petersen, 2016; Scimone et al., 2016). We used laminin and Hoechst staining to test double-RNAi animals for their ability to form a secondary or tertiary pharynx. *src-1* inhibition enhanced the penetrance of the ectopic pharynx phenotype after inhibition of *ndl-3* (from 11% in *ndl-3(RNAi)* to 40% in *ndl-3+src-1(RNAi)*) and *ptk7* (from 5% in *ptk7(RNAi)* to 58% in *ptk7+wntP-2(RNAi)*). Under these conditions, single-gene inhibition of *wntP-2* led to an already highly penetrant trunk duplication phenotype (88%), and while *src-1(RNAi)* did increase this slightly (94%), *src-*1+*wntP-2* RNAi caused a higher expressivity of forming two ectopic pharynges (Figure 5B). Together these results indicate the *src-1* inhibition sensitizes animals to disruption of either head or trunk control systems. The head and trunk PCG systems are thought to act independently, so we wondered whether *src-1* co-inhibition with PCGs might reveal hidden dependencies in the outputs to these systems. To test this, we inhibited *src-1* along with *ndl-3, ptk7,* or *wntP-2* and tested for effects on head patterning by counting *cintillo+* cells (Figure S5) and likewise inhibited *src-1* along with *ndk* and *wnt11-6* and tested for effects on trunk patterning by staining for *laminin* (Figure 5B). *src-1* co-inhibition with *ndl-3* mildly enhanced the increased brain cell number phenotype, but co-inhibition with *ptk7* and *wntP-2* had no additional effect on brain expansion (Figure S5). Therefore, after *src-1* inhibition, the activities of these trunk PCG regulators remained broadly tied to regulating trunk identity. Surprisingly however, co-inhibition of *src-1* along with PCGs that ordinarily only regulate head identity (*ndk, wnt11-6*) dramatically enhanced the penetrance of ectopic pharynx formation in regenerating trunk fragments (Figure 5B). Whereas 0% of control and 7% (1/13) of *src-1(RNAi)* animals formed extra pharynges, 75% (12/16) of *ndk+src-1(RNAi)* animals and 40% (6/15) of *wnt11-6+src-1(RNAi)* animals formed extra pharynges. By contrast, we have never observed that individually inhibiting *wnt11-6* or *ndk* leads to formation of ectopic pharynges. These results indicate that *src-1* inhibition can reprogram PCG outputs, implying that *src-1* ordinarily helps to channel regionalized factors into controlling the identity of specific anterior territories.

## Discussion

Together, these results suggest a distinct role for *src-1* in planarian regeneration in controlling anterior patterning (Figure 7). We found that *src-1* acts as a global negative regulator of anterior patterning, because its inhibition resulted in the expansion of both head and trunk identity. These effects were largely independent of hallmarks of head-tail AP axis polarization: injury-induced *wnt1* or *notum*, or the formation of *wnt1* and *notum* poles. Given that Src is an intracellular kinase known to act a downstream of multiple receptors (Erpel and Courtneidge, 1995; Thomas and Brugge, 1997; Abram and Courtneidge, 2000; Lemmon and Schlessinger, 2010), we sought to determine whether *src-1* could regulate anterior patterning downstream or in parallel to planarian Wnt and/or FGFRL signals.

Srcs have varied relationships to Wnt pathways described across several systems. In the canonical Wnt signaling pathway, Wnt binding to Frizzled receptors recruits Dishevelled (Dvl), sequestering Axin and preventing GSK3 phosphorylation of β-catenin that leads to its proteolysis through the destruction complex, thus allowing β-catenin accumulation and nuclear translocation to activate gene expression via TCF/LEF transcription factors (Gao and Chen, 2010). In mammalian F9 carcinoma cells, Src knockdown led to reduced canonical Wnt3a-stimulated TCF/LEF reporter output, an affect attributed to Src’s ability to bind and phosphorylate Dvl-2, potentiating activation of canonical Wnt downstream signals (Yokoyama and Malbon, 2009). Srcs can also act downstream of noncanonical Wnt pathways such as the Derailed/Ryk receptors transducing Wnt5 family signals important for neuronal development (Wouda et al., 2008; Petrova et al., 2013). Srcs can also function negatively in Wnt signaling, for example through targeting the Wnt co-receptor Lrp6/arrow for inactivation in developing zebrafish embryos (Chen et al., 2014). Src can also function downstream of the kinase-dead co-receptor Ptk7 (Andreeva et al., 2014), which acts as a Wnt co-receptor for either activating or inactivating canonical Wnt signaling (Peradziryi et al., 2011; Hayes et al., 2013). Together with these observations, the identification of a planarian Src that collaborates with Wnt-dependent processes supports a potentially ancient connection between these factors for axis formation.

We considered the possibility that *src-1* could act broadly downstream of several planarian Wnts. Some lines of evidence from this and prior work could be consistent with this interpretation. First, inhibition of planarian *dishevelled-2* caused the simultaneous formation of a secondary ectopic pharynx and posterior ectopic photoreceptors, similar to the *src-1(RNAi)* phenotype (Almuedo-Castillo et al., 2011). Second, the *src-1* RNAi phenotype resembled a combination of *wnt11-6* and *wntP-2* RNAi phenotypes and also caused a permanent shift to the site of eye regeneration similar to *wnt11-6* RNAi. While the apparent distinction between *src-1* and *beta-catenin-1* RNAi phenotypes suggests differences in their activities, the downstream factors in *wntP-2* and *wnt11-6* signaling are not fully understood, in part because of pleiotropic effects from *beta-catenin-1* inhibition. While reduced doses of *β-catenin-1* dsRNA have been reported to result in the formation of an ectopic posterior mouth and pharynx primordium (Almuedo-Castillo et al., 2011), the most prominent effects of *β-catenin-1* are highly penetrant formation of posterior and ectopic heads (Gurley et al., 2008; Petersen and Reddien, 2008; Adell et al., 2009). *src-1* appears to operate more primarily with the Wnt/FGFRL signals that pattern that AP axis versus the *wnt1*/*notum* pole signals responsible for determining head-versus-tail polarity. Therefore, it is unlikely that *src-1* transduces all Wnt signals in the animal or is involved in all instances of beta-catenin signaling.

However, our data is not consistent with a model in which *src-1* acts exclusively downstream of *notum,* and therefore its Wnt targets, to control anterior identity. Simultaneous inhibition of *src-1* and *notum* generated a synthetic phenotype in which animals displayed both phenotypes, as opposed to an outcome indicative of genetic epistasis (Figure 3A). In prior work, *wnt11-6* inhibition fully suppressed the *notum(RNAi)* ectopic eye phenotype, suggesting that *notum* primarily acts through *wnt11-6* for controlling eye placement (Hill and Petersen, 2015), and that *src-1* is unlikely to act primarily downstream of *wnt11-6*. However, we cannot rule out the possibility that *src-1* could act downstream of any Wnts that can act independently of *notum* and influence head regionalization. Testing this model would require future work to identify patterning roles for other negative regulators of specific Wnts or methods to detect Src activation. Interestingly, Dishevelled has also been shown to act in non-canonical Wnt signaling and to mediate a Wnt5-derailed/Related to tyrosine kinase (RYK)–dependent signal, which can signal through Src (Gao and Chen, 2010). Planarian *wnt5* defines the lateral-medial axis in planarian regeneration (Gurley et al., 2010), and inhibition of *dishevelled-1* in planarians has been shown to recapitulate aspects of the *wnt5(RNAi)* phenotype such as lateral separation of the planarian brain lobes (Almuedo-Castillo et al., 2011). However, *wnt5* is not believed to regulate anteroposterior patterning, so it is unlikely that *src-1* acts mainly downstream of *wnt5* to control AP head and trunk regionalization.

We also considered the possibility that *src-1* could act downstream of an unidentified receptor or FGFRLs. FGFRLs such as *ndk* and *ndl-3* have been shown to regulate regional identity in planarians (Cebria et al., 2002; Lander and Petersen, 2016; Scimone et al., 2016), but the mechanism by which this signaling occurs is unclear. FGFRLs have been shown to act as decoy receptors in *Xenopus* embryos. The FGFRL1 ectodomain is shed from the cell membrane and binds to some FGF ligands with high affinity, including FGF2, FGF3, FGF4, FGF8, FGF10, and FGF22 to regulate FGF signaling (Steinberg et al., 2010). However, inhibition of FGF or FGFRs in planarians have not resulted in patterning phenotypes. The intracellular domain of human FGFRL1 can interact with the Spred1 signaling molecule which could allow for downstream intracellular signaling (Zhuang et al., 2011). Furthermore, in beta-cell insulin granules, the intracellular domain of FGFRL1 can bind SHP-1 via a SH2 domain to activate ERK signaling (Silva et al., 2013). Given the synergist effects of *src-1* inhibition with *ndk* and *ndl-3* (Figure 5A-D) and the similar synergistic RNAi phenotypes seen with *ndk* and *fzd5/8-4* (Scimone et al., 2016) it was possible that cross-regulation between WNT and FGFRL signaling pathways controls body regionalization in planarians and that this may be mediated through *src-1*. However, the examination of the extra photoreceptor defect in *src-1* RNAi revealed more similarities with the *wnt11-6* phenotype than the *ndk* phenotype. First, *ndk* RNAi generated posterior photoreceptors located more distantly than either *wnt11-6* or *src-1* RNAi. Second, both *wnt11-6* and *src-1* RNAi tended to shift the location of eye regeneration more posteriorly, unlike *ndk* RNAi. These observations suggest *src-1* acts distinctly from *ndk*. Other signaling factors known to interface with Src have been described in planarians but are unlikely to explain the patterning roles of *src-1*. For example, integrins are well known to signal through Src for adhesion and signaling (Huttenlocher and Horwitz, 2011) but inhibition of the single planarian integrin-beta led to tissue disorganization and excess neurogenesis in general rather than specific AP patterning effects seen in *src-1* RNAi (Bonar and Petersen, 2017; Seebeck et al., 2017), but it remains possible that AP patterning involves some input from integrin signals. Future systematic analysis of the many possible upstream Src receptors could help resolve the role of Src for AP patterning with respect to Wnts. Thus, taken together, we propose a role for *src-1* in globally suppressing anterior identity and regulating posterior determination in parallel to the action of Wnts and FGFRLs, perhaps using alternative signal inputs.

Our analysis of dual inhibition phenotypes between *src-1* and known AP patterning regulators further suggests this model of parallel action. The expressivity of any patterning phenotype we examined that involved anteriorization of posterior tissue was enhanced dramatically after *src-1* RNAi, which would be expected if *src-1* and Wnts act independently to control pattern. Other explanations are possible, for example that *src-1* acts downstream of all Wnt genes and that the relatively weaker Wnt RNAi phenotypes represent incomplete knockdown. The ability for *src-1* RNAi to reprogram the outputs of anterior PCGs into controlling trunk identity suggests *src-1* may act in a buffering process that helps channel PCG factors into controlling distinct outputs. In addition, the progressive nature of the *src-1* RNAi phenotype to affect anterior regions more strongly than posterior regions was consistent with the finding that PCG domains were shifted rather than eliminated by *src-1* RNAi. The outcomes of double-RNAi between *src-1* and head or trunk PCGs are also suggestive of an anterior bias to *src-1* function in which, at least in within the context of *src-1* inhibition, anterior factors like *ndk* can be reprogrammed to influence more posterior identity but more posterior factors like *ptk7* and *wntP-2* are not reprogrammed to influence more anterior identity. This could ultimately reflect the overlapping uses of the two Wnt/FGFRL systems to define non-anterior in successive domains. It is also possible that the hypothesized parallel actions of *src-1* and Wnts could arise from distinct signaling to regulate PCG domains within the muscle versus interpretation of domain identity via neoblast-dependent tissue formation. Reagents to examine Src activation status, along with systematic tests of the many possible receptors upstream of Src, will be helpful in resolving these and other mechanisms. It is intriguing to note that the Src family kinase SRC-1 in *C. elegans* acts in parallel to Wnt signaling in order to regulate spindle orientation along the AP axis of the very early embryo (Bei et al., 2002). These observations suggest there may be deep ancestry to the use of joint activities for Src family kinases and Wnt signals in forming the primary body axis. Together our results identify *src-1* as a new factor regulating positional information along the planarian AP axis that is used for specifying the proper identity of missing tissues in regeneration.

## Materials and Methods

### Planarian culture

Asexual strain CIW4 of the planarian *Schmidtea mediterranea* were maintained in 1× Montjuic salts at 19°C as described (Petersen and Reddien, 2011). Planarians were fed a liver paste and starved for at least 7 days before experiments.

### Cloning

*src-1* (dd_Smed_v6_3147_0_1) was identified through blast searching the planarian transcriptome at https://planmine.mpibpc.mpg.de/ (Brandl et al., 2015; Brandl et al., 2016))(Grohme et al., 2018). Primers used for cloning *src-1* were 5′-AAGCTTGGTGGCTTGCTTTA-3′ and 5′-TGCGATCAACCAATGAAAAA-3′. Primers for cloning the genes from the screen are indicated in Table S1.

### Riboprobes

Riboprobes and double-stranded RNA (dsRNA) for *src-1* were generated by *in vitro* transcription (NxGen, Lucigen) as described previously (Petersen and Reddien, 2011). Riboprobes and dsRNAs for *src-1* were cloned by RTPCR into pGEM-T-easy using the following primers: *src-1:* 5′-AAGCTTGGTGGCTTGCTTTA-3′ and 5′-TGCGATCAACCAATGAAAAA-3′ Other riboprobes (*chat, cintillo, gluR, opsin, tyrosinase, collagen, laminin, notum, wnt1, ndk, ndl-5, fzd4, wntP-2, axinB, ptk7*) were as previously described (Oviedo et al., 2003; Cebria et al., 2007; Reddien et al., 2007; Wang et al., 2007; Collins III et al., 2010; Gurley et al., 2010; Wenemoser and Reddien, 2010; Petersen and Reddien, 2011; Lapan and Reddien, 2012; Currie and Pearson, 2013; März et al., 2013; Vu et al., 2015)

### RNAi

RNAi was performed either by dsRNA feeding. For RNAi, dsRNA was synthesized from *in vitro* transcription reactions (NxGen, Lucigen). dsRNA corresponding to *Caenorhabditis elegans unc-22*, not present in the planarian genome, served as a negative control. For the RNAi screen (Table S1), animals were fed a mixture of liver paste and dsRNA three times over 6 days, then amputated transversely to generate head, trunk, and tail fragments. Animals were scored for regeneration defects after 14 days of regeneration (Table S1). For other experiments unless noted otherwise, animals were fed a mixture of liver paste and dsRNA six times in 14 days prior to amputation of heads and tails 4 h after the final feeding. For Figure 4, animals were fed dsRNA 12 times over 6 weeks and starved for 3 days before eye resection. For all comparisons between double RNAi and single RNAi conditions, an equal amount of control competing dsRNA was mixed with the single RNAi condition so that animals across treatments received the same overall amount of dsRNA.

### In situ hybridization and immunostaining

Colorimetric (NBT/BCIP) or fluorescence *in situ* hybridizations were performed as described (Lander and Petersen, 2016) after fixation in 4% formaldehyde and bleaching (Pearson et al., 2009) using blocking solution containing 10% horse serum and western blot blocking reagent (Roche) (King and Newmark, 2013). Digoxigenin- or fluorescein-labeled riboprobes were synthesized as described (Pearson et al., 2009) and detected with anti-digoxigenin-HRP (1:2000, Roche/Sigma-Aldrich 11207733910, lot 10520200 anti-fluorescein-HRP (1:2000, Roche/Sigma-Aldrich 11426346910, lot 11211620) or anti-digoxigenin-AP (1:4000, Roche/Sigma-Aldrich 11093274910, lot 11265026). Hoechst 33342 (Invitrogen) was used at 1:1000 as a counterstain. For immunostainings, animals were fixed in Carnoy’s solution as described (Hill and Petersen, 2015), using tyramide amplification to detect labeling with rabbit anti-6G10 (1:3000, Cell Signaling D2C8, lot 3377S).

### Image analysis

Live animals and NBT/BCIP-stained animals were imaged with a Leica M210F dissecting microscope and a Leica DFC295, with adjustments to brightness and contrast using Adobe Photoshop. Whole animal fluorescence imaging was performed on either a Leica DM5500B compound microscope with Optigrid structured illumination system or a Leica laser scanning SPE confocal microscope at 40× or 63×, and presented images are maximum projections of a *Z*-series with adjustments to brightness and contrast using ImageJ and Photoshop. Plots were generated in Microsoft Excel or R (ggplot2).

### Cell counting

*cintillo^+^* cells in the brain were counted manually and normalized to the square root of the animal area determined using Hoechst staining and CellProfiler (Lamprecht et al., 2007).

### Real-time PCR

Total RNA was extracted by mechanical homogenization in Trizol (Life Technologies), DNase-treated (TURBO DNAse, Ambion), and reverse transcribed with oligo-dT primers (Multiscribe reverse transcriptase, Applied Biosystems), and qPCR was performed using Eva Green PCR Master Mix (Biotium) from nine regenerating fragments in four (Figure 4.2) biological replicates. Relative mRNA abundance was calculated using the delta-Ct method after verification of primer amplification efficiency, normalizing to ubiquilin expression. *P*-values below 0.05 by a two-tailed *t*-test were considered as significant. The following primer sets were used: *src-1:* 5′-ATGACGTGTATAACGCCGACAC-3′, 5′-TGAGGACAGGACAGTGTTAATTTG-3′ *ubiquilin:* 5’-ATTCGTCGGAATTGGAAACA-3’, 5’-GCGTTCACATCTCCAAAGGT-3’

### Eye Regeneration Assays

Modified from (Hill and Petersen, 2018). Briefly, worms were immobilized on ice for resection. Eyes were removed using a hypodermic needle. All animals were tracked individually and imaged one day prior to eye removal, one day after eye removal to confirm resection of eye tissue, and 22 days post-surgery to determine the regenerative outcome.

## Acknowledgements

We thank members of the Petersen lab for critical comments. C.P.P. acknowledges funding from the National Institutes of Health, USA (NIGMS R01GM129339 and R01GM130835), and pilot project funding from the NSF-Simons Center for Quantitative Biology at Northwestern University, an NSF (1764421)-Simons/SFARI (597491-RWC) MathBioSys Research Center.

## Author Contributions

C.P.P. conceived of the study and acquired funding, N.B. and D.G. designed and conducted experiments, and C.P.P., N.B. and D.G. wrote the manuscript.

## Supplementary Figures

**Supplementary Figure 1.**
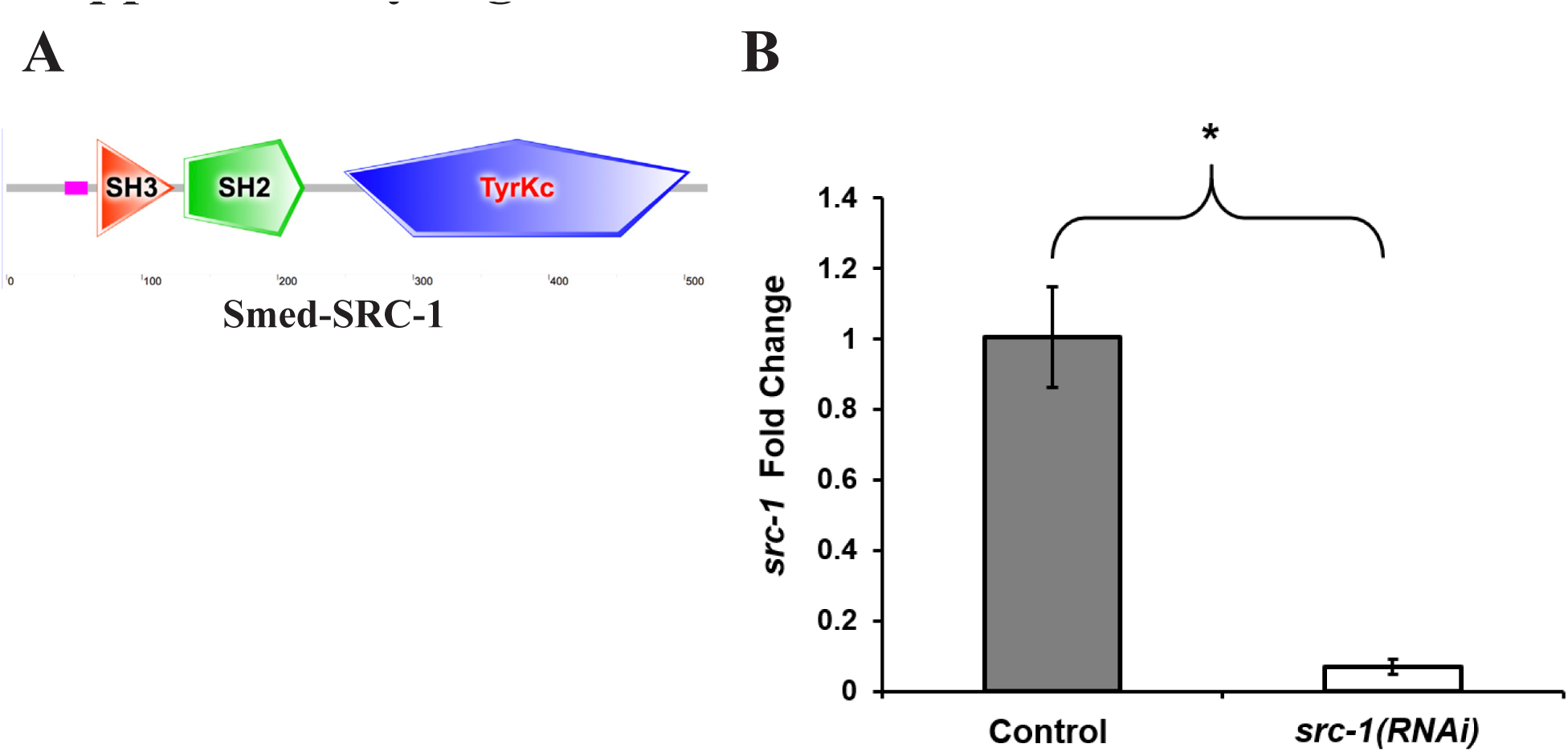
(A) Domain structure of dd_Smed_v6_3147_0_1 (*src-1*) contains an SH3 (E-value = 3.07e-17), SH2 (E-value= 4.73e-20), and Tyrosine Kinase domain (E-value 9.29e-124) indicative of Src- family kinases. (B) qPCR to detect RNAi knockdown of *src-1* mRNA after treatment of animals with *src-1* dsRNA versus control dsRNA. * p<0.05, two-tailed t-test.

**Supplementary Figure 2.**
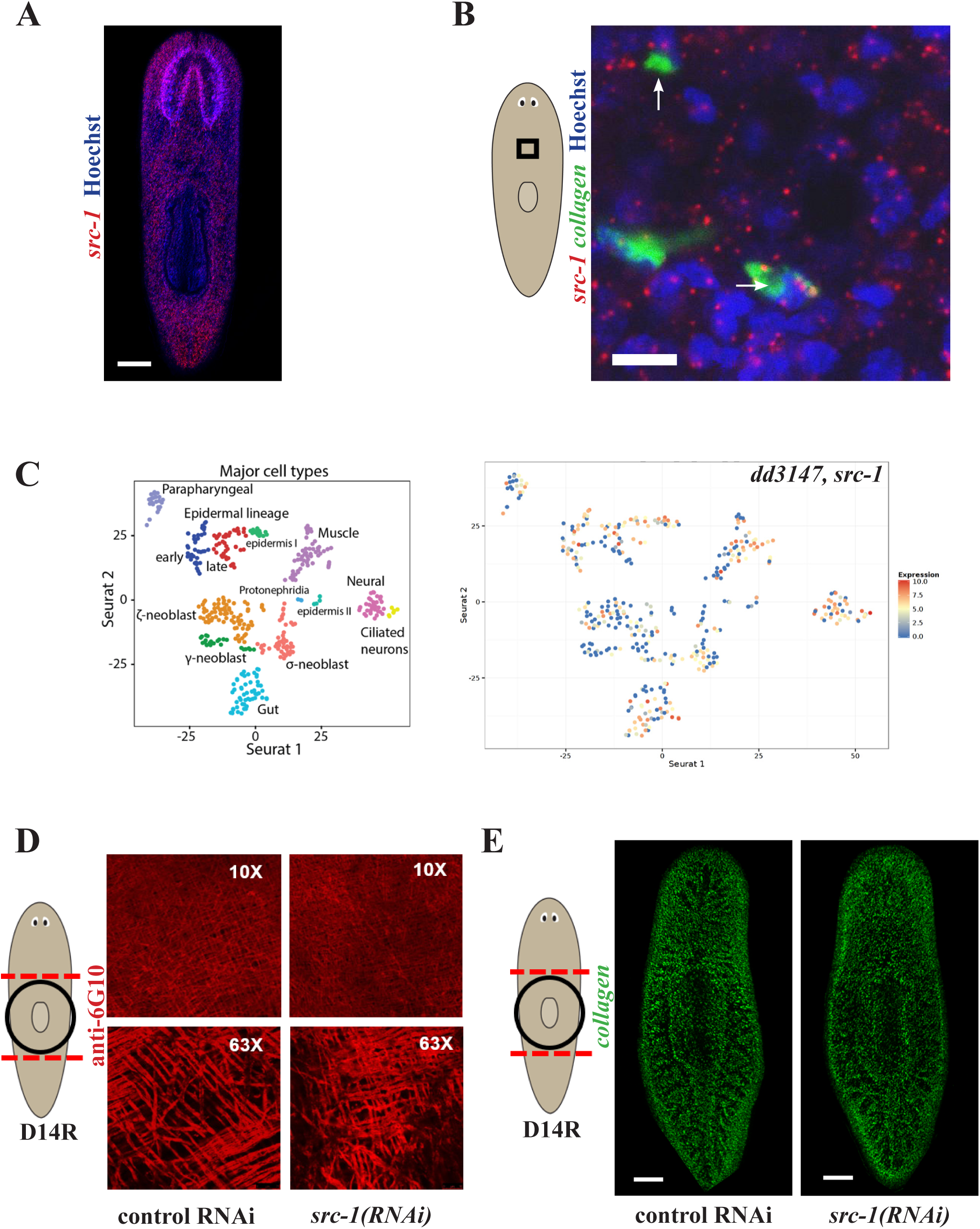
(A) FISH to detect *src-1* expression, showing broad expression throughout the planarian body. Scale bars, 150µm. (B) Double FISH to detect the expression of *src-1* and *collagen* in uninjured animals, with cartoons indicating the approximate location of the imaged regions. *src-1* mRNA expression was punctate and broad. Some *collagen*+ cells could be identified with overlapping detection of *src-1* FISH signal (arrows). (C) Single-cell RNA-seq expression profiling as measured from a prior study (Wurtzel et al., 2015) detected *src-1* transcripts in epidermis, muscle, gut, neurons, neoblasts, and other cell types. (D) Muscle fibers (anti-6G10) were present in *src-1(RNAi)* regenerating animals. (E) Muscle cell bodies (FISH for *collagen* expression) were present in *src-1(RNAi)* regenerating animals.

**Supplementary Figure 3.**
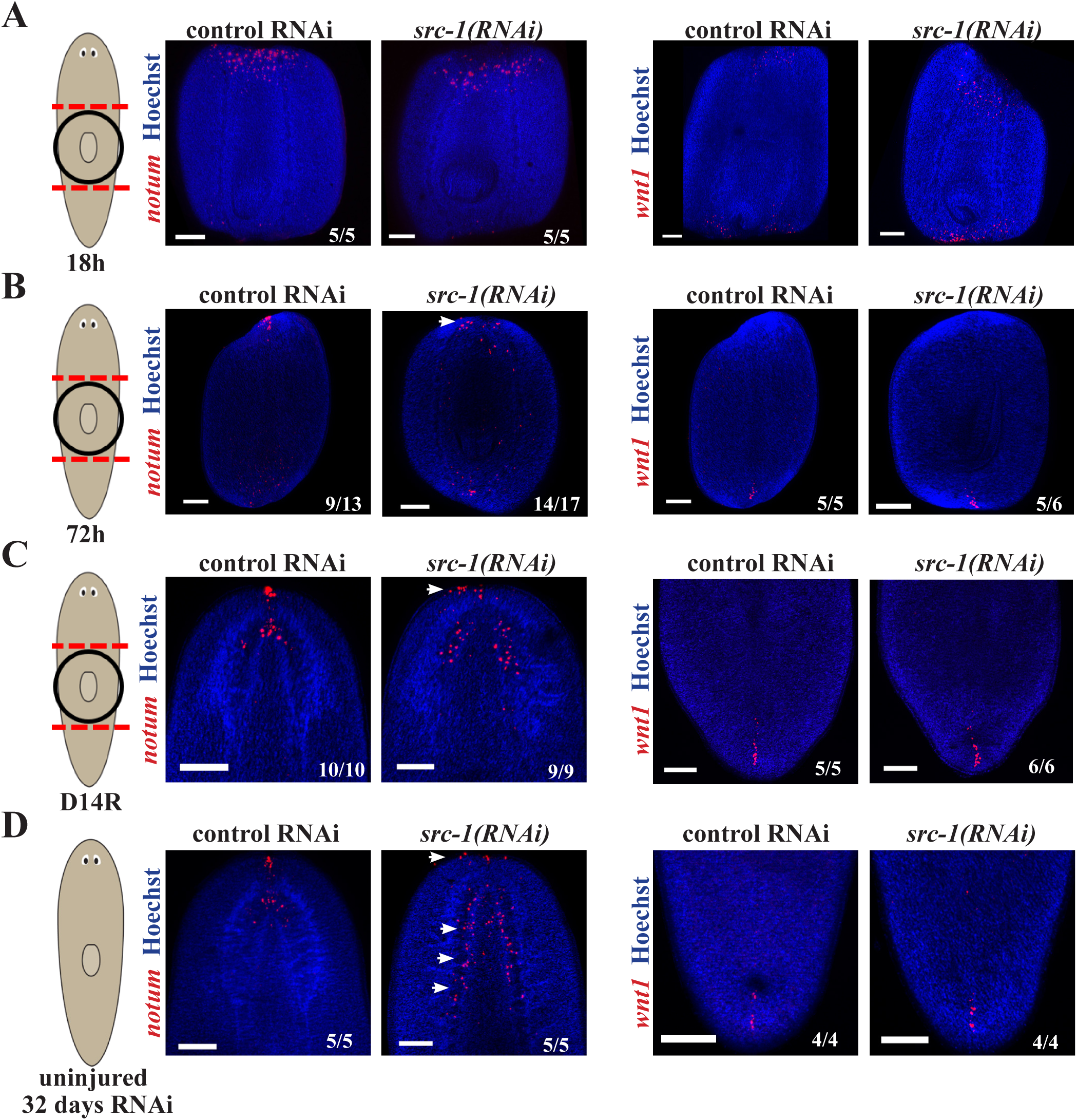
Expression of polarity determinants *notum* and *wnt1* in *src-1(RNAi)* animals (A) 18 hours after amputation, (B) 72-hours after amputation, (C) after 14 days of regeneration, and (D) after 3 weeks of homeostatic knockdown of *src-1*. (A) *src-1(RNAi)* trunk fragments have normal wound-induced *notum* (left) and *wnt1* (right) expression at 18 hours post amputation. (B) *src-1(RNAi)* trunk fragments at 72-hours post-amputation show a delay in formation of the *notum-*expressing anterior pole (14/17 animals showed disperse notum expression and 3/17 showed focused expression) compared to control animals (9/13 had focused expression by 72 hours and 4/13 had disperse expression). Right, *src-1(RNAi)* animals underwent normal formation of the *wnt1-*expressing posterior pole (C) *src-1(RNAi)* animals regenerating their head ultimately form an anterior expanded anterior pole and also expanded domain of brain-associated *notum* in conjunction with their overall expanded brain at 14-days post-amputation (arrows) but normal *wnt1* expression. (D) Uninjured *src-1(RNAi)* animals have a slightly expanded anterior pole and more brain *notum* but normal *wnt1* expression. Scale bars, 150 µm.

**Supplementary Figure 4.**
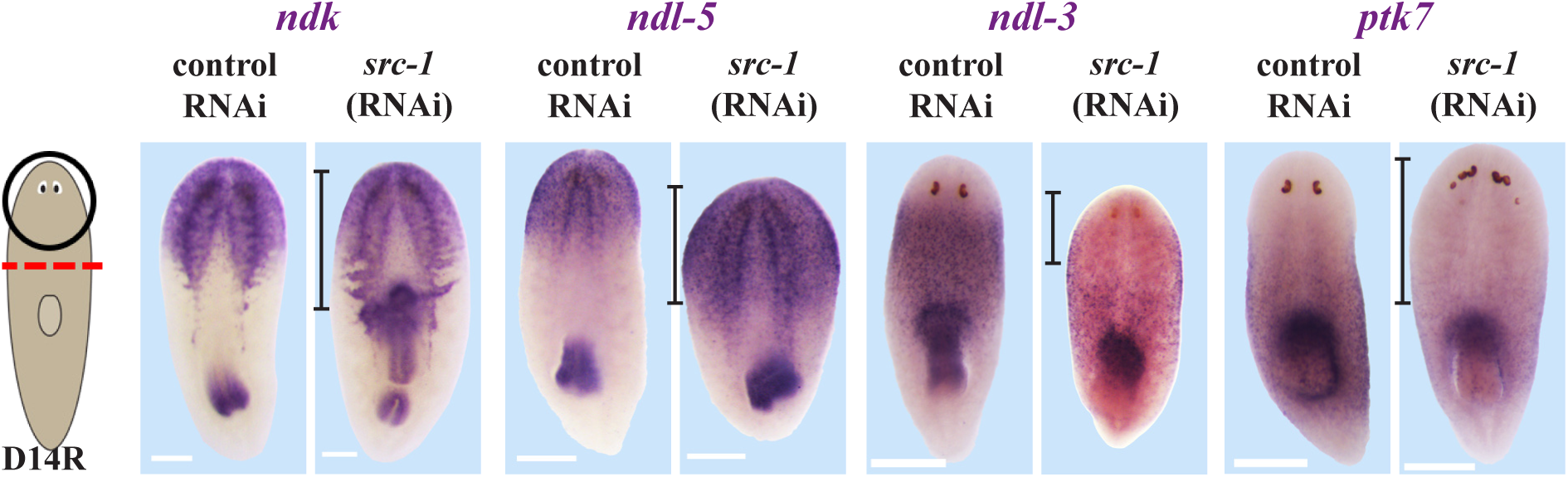
*src-1(RNAi)* versus control RNAi regenerating head fragments fixed at 14 days post amputation and stained by WISH for expression of anterior PCGs as indicated, panels represent at least 5/5 animals as shown.

**Supplementary Figure 5.**
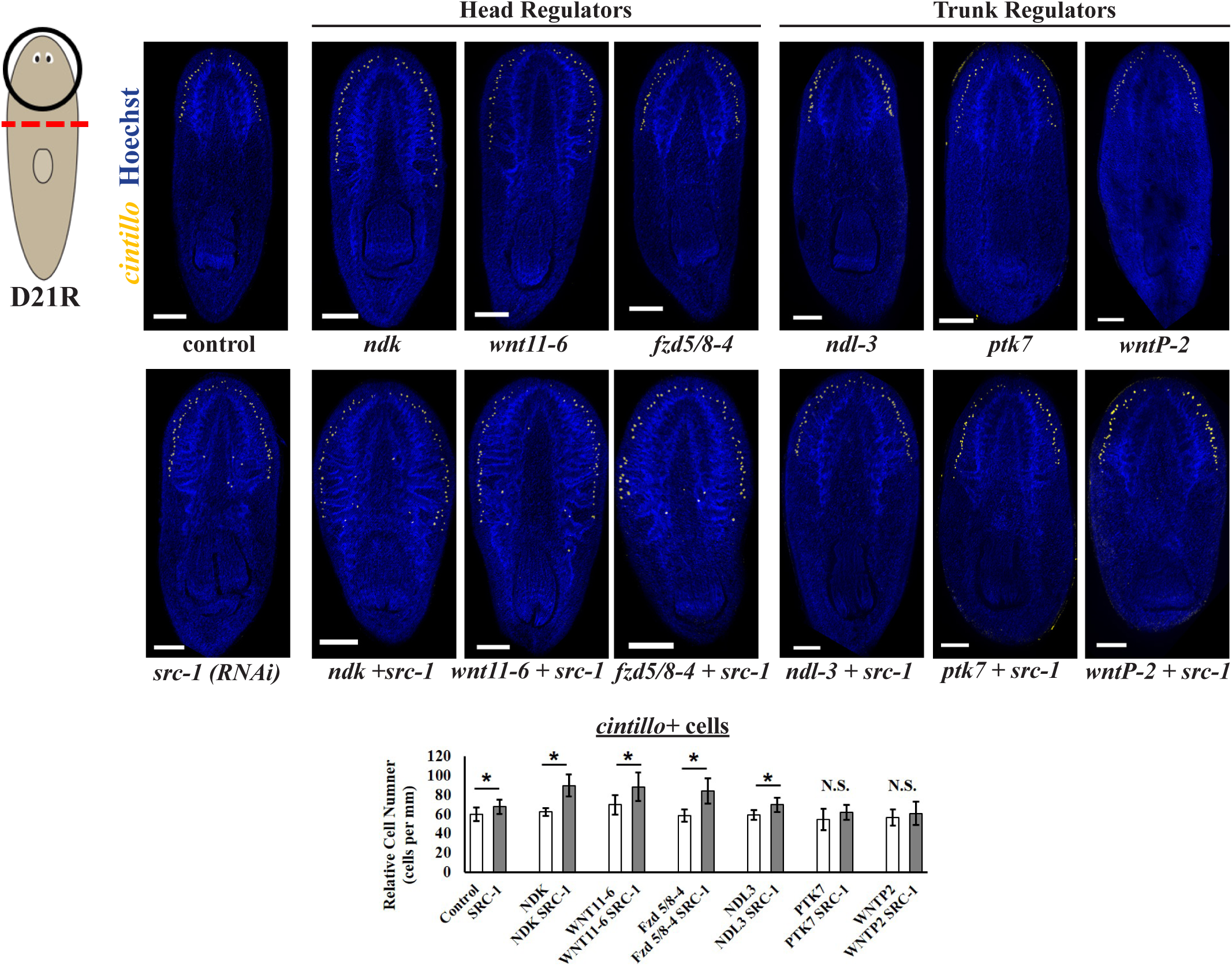
(A) FISH to detect expression of *cintillo* (yellow), a marker of chemosensory neurons, in head fragments at day 21 post amputation. Hoechst (blue) used as counterstain to detect nuclei. Simultaneous inhibition of *src-1* with *wnt11-6*, *ndk*, or *fzd5/8-4* resulted in the formation of a larger brain as determined by the greater relative number of *cintillo+* cells compared to single-gene inhibitions of *src-1*, *wnt11-6*, *ndk*, or *fzd5/8-4* alone. Simultaneous inhibition of *src-1* with patterning factors known to restrict trunk identity, *ptk7* or *wntP-2*, did not result in the formation of a larger brain greater than the effects *src-1(RNAi)* alone. *ndl-3+src-1* RNAi weakly increased numbers of *cintillo+* cells as measured. Scale bars, 150um. Graph shows quantification of *cintillo+* cell number normalized to animal size. *, p <0.05 from 2-tailed t-test.

